# Species-Specific Root Microbiota Dynamics in Response to Plant-Available Phosphorus

**DOI:** 10.1101/400119

**Authors:** Natacha Bodenhausen, Vincent Somerville, Alessandro Desirò, Jean-Claude Walser, Lorenzo Borghi, Marcel G.A. van der Heijden, Klaus Schlaeppi

**Author notes:** Corresponding author: Klaus Schlaeppi, University of Bern, Institute of Plant Sciences, Altenbergrain 21, 3013 Bern.

## Abstract

- Phosphorus (P) is a limiting element for plant growth. Several root microbes, including arbuscular mycorrhizal fungi (AMF), have the capacity to improve plant nutrition and their abundance is known to depend on P fertility. However, how complex root-associated bacterial and fungal communities respond to changes in P availability remains ill-defined.
- We manipulated the availability of soil P in pots and compared the root microbiota of non-mycorrhizal Arabidopsis with mycorrhizal Petunia plants. Root bacteria and fungi were profiled using ribosomal operon gene fragment sequencing, we searched for P sensitive microbes and tested whether a P sensitive core microbiome could be identified.
- Root microbiota composition varied substantially by P availability. A P sensitive core microbiome was not identified as different bacterial and fungal groups responded to low-P conditions in Arabidopsis and Petunia. P sensitive microbes included Mortierellomycotina in Arabidopsis, while these were AMF and their symbiotic endobacteria in Petunia. Of note, their P-dependent root colonization was reliably quantified by sequencing.
- The species-specific root microbiota dynamics suggest that Arabidopsis and Petunia evolved different microbial associations under the selection pressure of low P availability. This implies that the development of microbial products that improve P availability requires the consideration of host-species specificity.

## Introduction

Phosphorus (P) presents one of the key nutrients for plant growth. While a small fraction of soil P is directly available for plant uptake, the larger fraction is complexed to organic and mineral soil components and therefore inaccessible for plants. The conventional agronomic solution to increase P availability for plants relies on supplementing mineral phosphate (PO_4_^3-^). However, yield optimization requires excess application of phosphate since less than thirty percent of applied P fertilizers effectively support plant growth, the rest of the applied phosphate readily transforms to plant non-available P forms (Cordell *et al.*, 2009). Such an overuse causes the rapid depletion of finite phosphorus reservoirs, is expensive and causes environmental harm, primarily with negative impacts on the aquatic environment by eutrophication of the surface water (Cordell *et al.*, 2009; Scholz & Wellmer, 2013; Reijnders, 2014). Therefore, next-generation agriculture requires novel sustainable solutions that reduce fertilizer inputs and increase the nutrient-use efficiency while maintaining high plant yields.

Numerous root-associated microbes have the capacity to mobilize soil P or metabolize recalcitrant forms (Gyaneshwar *et al.*, 2002; Jacoby *et al.*, 2017), thereby enhancing plant performance and agricultural yield especially under nutrient limiting conditions. For fungi, there is a continuum of functionally similar associations between different groups of root fungi and their host plant species (van der Heijden *et al.*, 2017). For instance, arbuscular mycorrhizal or ectomycorrhizal fungi are intimately connected to plant roots and support plant growth by mobilizing and transporting P from a larger soil volume and more distant pools of P thanks to their large hyphal network, (Jakobsen *et al.*, 1992). Aside from these classical mycorrhizal plants, also nonmycorrhizal plants such as *Arabidopsis thaliana* [hereafter: Arabidopsis] and *Arabis alpina* rely on fungal associations for nutrient acquisition (Cosme *et al.*, 2018). They rely on beneficial fungal endophytes including *Colletotrichum tofieldiae* (Hiruma *et al.*, 2016), *Serendipita indica* [formerly *Piriformospora indica*] (Yadav *et al.*, 2010) or a fungus of the order Helotiales (Almario *et al.*, 2017). Typically, the P availability in soil determines to which extent a plant is colonized by the fungal symbiotic partner with high levels of colonization under low-P conditions and little colonization in soils with high-P levels. In addition to fungi, many root bacteria are known to support plant nutrition with their abilities to solubilize inorganic P or to mineralize organic P (Rodríguez & Fraga, 1999; Alori *et al.*, 2017). Powerful P solubilizing bacteria include strains from the genera *Bacillus, Pseudomonas* and *Rhizobium*; for a more comprehensive list as well as their growth effects on crops, we refer to Alori *et al*. (2017). While a wide range of individual rhizosphere microbes is known to support plant P nutrition, the effects of P availability on the overall root microbiota remains less understood. A deeper understanding of interactions between plants and their microbial allies in response to the bioavailability of P is needed for developing microbe-dependent P fertilization solutions (Schlaeppi & Bulgarelli, 2015; Busby *et al.*, 2017).

Interactions among microbes emerge as a critical component for the maintenance of host-microbial homeostasis and for plant performance (Hassani *et al.*, 2018). Inter-kingdom microbial associations occur in the plant root microbiota as for instance root fungi hosting endobacteria in their cells (Desirò *et al.*, 2014). Such ancient fungi-endobacteria interactions (Bonfante & Desirò, 2017) include root fungi of the Mucoromycota (Spatafora *et al.*, 2017) that host diverse bacterial endosymbionts related to *Burkholderia* or *Mycoplasma*. An example of *Burkholderia*-related endobacteria includes *Candidatus* Glomeribacter gigasporarum that is hosted by a Glomeromycotina fungus (Bianciotto *et al.*, 2003). *Mycoplasma*-related endobacteria have a broader host range with presence in Glomeromycotina (Naumann *et al*., 2010), Mortierellomycotina (Desirò *et al*., 2018) and Mucoromycotina (Desirò *et al*., 2015). The occurrence and functional contribution of fungal endobacteria adds a further level of complexity to the interactions of plant with and among their associated microbes.

Plants are more and more recognized in context with their microbial communities, where a multitude of microbes collectively function as a microbiome. P fertilization as well as P depletion are known to induce shifts in soil microbial communities (Wakelin *et al.*, 2012; Leff *et al.*, 2015; Huang *et al.*, 2016; Bergkemper *et al.*, 2016; Ikoyi *et al.*, 2018). For example, grassland soil microbes consistently responded to phosphate inputs with compositional community changes, as for instance mycorrhizal fungi, oligotrophic bacteria and methanogenic Archaea decreased in relative abundance with nutrient additions (Leff *et al.*, 2015). While the responses of soil microbial communities to varying levels of different sources of P have been well studied, the plant root-associated microbial communities have received less attention (Silva *et al.*, 2017; Almario *et al.*, 2017; Robbins *et al.*, 2018). Robbins *et al*., (2018) investigated the effects of different levels of P fertilization on the Arabidopsis rhizosphere and root microbiota. While phosphate applications had little effects on microbial diversity, they affected more strongly the plant-associated microbiota compared to bulk soil communities, suggesting plant-mediated cues for structuring the plant microbiota in response to the nutritional status. The authors noted a weak P-fertilization effect on root communities that was manifested by low-abundant root-associated microbes. This suggests P to be a minimal driver in shaping microbial communities compared to larger drivers such as compartment (soil vs. rhizosphere, rhizoplane, and roots) or soil type (soils differing chemically, physically and with regard to their microbiota; Hacquard *et al.*, 2015). The work on model non-mycorrhizal plants revealed subtle responses of the root microbiota to the availability of soil P; further work is needed to test whether mycorrhizal plants exhibit stronger root microbiota responses thanks to their colonizing symbionts.

*Petunia x hybrida* [hereafter: Petunia] is a model plant that is commonly used for investigating the symbiosis with arbuscular mycorrhizal fungi (AMF, Wegmüller *et al.*, 2008; Breuillin *et al.*, 2010). Due to its fast life cycle, modest size and the availability of genetic tools made, Petunia is also a model to study plant development (Vandenbussche *et al.*, 2016), plant nutrition (Liu *et al.*, 2018) and hormonal signaling (Hamiaux *et al.*, 2012). Petunia belongs to the Solanaceae family, thus is related to tomato, potato and eggplant so that root microbiota knowledge may be transferable to these staple food crops. We therefore chose Petunia in comparison with non-mycorrhizal Arabidopsis to study root microbiota dynamics in response to P availability in soil and how plants cope with P limiting conditions.

In this study, we tested the hypothesis that the composition of the root microbiota alters depending on the P availability and we asked whether a core microbiome or plant species-specific microbiomes prevail in response to P deficiency. We expected the Petunia root microbiota to enrich for AMF under low-P conditions, whereas the Arabidopsis response to low-P remained unclear. Hence, while differential fungal responses were anticipated for the two plant species, we were interested in their bacterial responses and whether a different sets of bacteria will respond to the varying levels of P availability. A particular goal of the study was to uncover the interplay between root bacteria and fungi and we examined their co-occurrence patterns in response to the varying P availability. We expected to find potential microbial interactions and hypothesized that the root microbiota data contains paired sequence information of fungal endobacteria and their corresponding host fungi. A technical goal of this study was to quantify AMF colonization in the context of whole fungal diversity based on DNA-based sequencing instead of the traditional morphological quantification by microscopy.

This study reveals that root microbiota composition varies markedly by P availability in soil and that different fungal and bacterial groups are responsive to low-P conditions in Arabidopsis and Petunia. We find co-abundant groups of candidate microbial cooperation partners, including AMF and their symbiotic endobacteria, both known to support plant growth under low-P conditions. Our work suggests that Arabidopsis and Petunia have evolved individual microbial solutions, involving multitrophic microbial interactions, to cope with low-P conditions.

## Material and Methods

### Plant Growth

The experiment was conducted in 400 ml pots lined with a mesh (Trenn-Vlies, Windhager, Thalgau, Germany). Soil was collected on April 4^th^ 2014 from a field site (47°26′20″ N 8°31′40″ E), sieved to 2 mm and stored at 4°C until use. Soil was mixed 1:1 volume with sterilized quartz-sand. Chemical properties of the sand-soil mixture were analyzed at the Labor für Boden-und Umweltanalytik (Eric Schweizer AG, Thun, Switzerland): pH 6.8, 6/31/51% (clay/silt/sand) and 1.4/1.05/1.07 mg kg-1 (water-extractable N/P/K).

Petunia seeds were surface sterilized with 70% ethanol, washed with autoclaved water and plated on 1/2 strength Murashige and Skoog basal medium (Sigma, Buchs, Switzerland) supplemented with 1.5% sucrose and solidified with 1.5% agar. Plants were germinated under long-day conditions (16-h photoperiod) in climate chamber (Sanyo MLR-352H; Panasonic, Osaka, Japan) at 25°C and 60% relative humidity. After 7 days, seedlings were transferred to 400 ml pots filled with substrate. Plants were grown for two weeks in the same climate chamber then moved to an in-house climate chamber with same humidity, photoperiod and temperature. Plants were fertilized with a corrected Petunia nutrient solution (Reddy *et al.*, 2007), prepared with three concentrations of phosphate: 0.03 mM KH_2_PO_4_ (low-P), 1 mM KH_2_PO_4_ (medium-P) and 5 mM KH_2_PO_4_ (high-P). Each plant received 300 ml of the solution over the last six weeks before harvest. We conducted two separate experiments using the same treatments and growth conditions, the first to collect the plant root samples (DNA analyses and microscopy) and shoot biomass and a second experiment to quantify leaf nutrient levels.

### Sample Collection

Plants were harvested at 10 weeks. The roots were separated from the shoot with a clean scalpel. The shoots were dried in a 60° C oven for dry weight analysis. The loosely attached soil was shaken from the roots, the roots were washed three times in PBS buffer (approximately 10 ml for 1 g of fresh weight) and then split into two equivalent subsamples. Samples for DNA extraction were stored at −80°C until processing. Samples for microscopy were stored in 50% ethanol. After staining with pen ink (Vierheilig *et al.*, 1998), root length colonization was determined using the magnified intersections method for 100 intersections per sample (McGonigle *et al.*, 1990). Soil from unplanted pots was collected by removing the top 1 cm layer and then mixing the soil below, one sample (250 mg - 500 mg) was taken from each pot. The dried shoots were weighed and milled. P and K concentrations were analyzed using inductively coupled plasma-optical emission spectroscopy (ICP-OES) at the elemental analytic department of Agroscope according to (VDLUFA-Verlag, 2006).

### Microbiota Profiling

The protocol for microbiota profiling, including DNA extraction, PCR, sequencing and bioinformatics, is described in detail in **Methods S1**. The comparison of the PCR approaches is reported in **Notes S1**, which contains the bioinformatic script, input data, analysis script and the markdown report. The bioinformatic analysis of the main samples of the study is documented with the scripts, parameters, support and report files in **Notes S2**. The raw sequencing data of the MiSeq runs and the SMRT sequencing are available from the European Nucleotide Archive ENA under the study accession PRJEB27162.

### Statistical Analyses

Statistical analyses were performed using R v3.3.2 (R Core Team, 2016) within Rstudio (RStudio Team, 2015). The effects of P availability on dry weight, P content and K content were assessed with a linear model. Dry weight data was log-transformed to satisfy the assumptions of the linear model (normality of residuals and homoscedasticity). To test for the effect of P availability on AMF colonization, a generalized linear model was fitted with quasibinomial distribution to account for overdispersion. Rarefaction curves were prepared with the function ‘rarecurve’ from vegan (Oksanen *et al.*, 2018). For alpha diversity, the data was rarefied to 15’000 sequences 500 times. For each subsample, several diversity indices were estimated: richness (S) is the number of OTUs, H is the Shannon index from which D=exp(H) was calculated (Jost, 2007), and Sheldon evenness is E= exp(H)/S (Sheldon, 1969). ANOVA was used to assess the effect of P a and plant species on the mean of the 500 subsamples for each sample. For the rest of the analysis, the data was filtered (at least 4 sequences per sample in 4 samples) to remove low abundant OTUs. The effects of P availability and plant species on community composition were assessed with permutational multivariate analysis of variance (PERMANOVA) of Bray-Curtis dissimilarities and visualized with principal coordinate analysis (PCoA) using vegan and phyloseq (McMurdie & Holmes, 2013). The effect of P availability on abundance of each OTU was investigated with edgeR (Robinson *et al.*, 2010) on TMM-normalized data (Robinson & Oshlack, 2010) and visualized with ternary plots. TMM-normalized data was used to calculate Spearman rank correlations between OTUs for co-occurrence networks. Positive (ρ>0.7) and significant relationships (*P*<0.001) were visualized with igraph (Csardi & Nepusz, 2006). Scripts, functions and support files are available as **Notes S3**. **Figure S1** visualizes the workflow of the analysis steps.

### Identification of Endobacteria

We describe the identification of endobacteria OTUs using a phylogenetic placement approach in **Methods S1**. Briefly, we pre-selected candidates in the microbiome dataset using two approaches and then validated their representative sequences by fine mapping to a reference tree of known endobacteria sequences. The first approach was based on sequence clustering and for the second, we employed co-occurrence characteristics from network analysis. Command line and analysis code in R (including markdown report) as well as the database with curated endobacteria 16S rDNA reference sequences are available as **Notes S4**.

## Results

### Experimental Setup for Manipulating Phosphate Levels

We investigated the dynamics of the root-associated microbiota to the availability of soil P and compared the non-mycorrhizal model species Arabidopsis to Petunia, which forms symbiosis with AMF. Plants were sown in a field soil that was amended with sand and we manipulated soil P availability by applying low, medium or high levels of phosphate. Unplanted pots were included as controls to collect soil samples. We first confirmed the effectiveness of the applied phosphate levels and found that phosphate treatments positively affected plant growth (**Fig. 1a**), increased P levels in plant leaves (**Fig. 1b**), while reducing the AMF colonization levels in Petunia roots (**Fig. 1c**). As we manipulated soil P levels using simple K salts, we tested if the low-P condition would also be limited in potassium. Since the plants growing in low-P conditions were sufficiently supplied with K (**Fig. 1d**), we concluded that the simple approach of using KH_2_PO_4_^3-^ solutions permitted to establish a P gradient without causing K limiting conditions.

**Figure 1:**
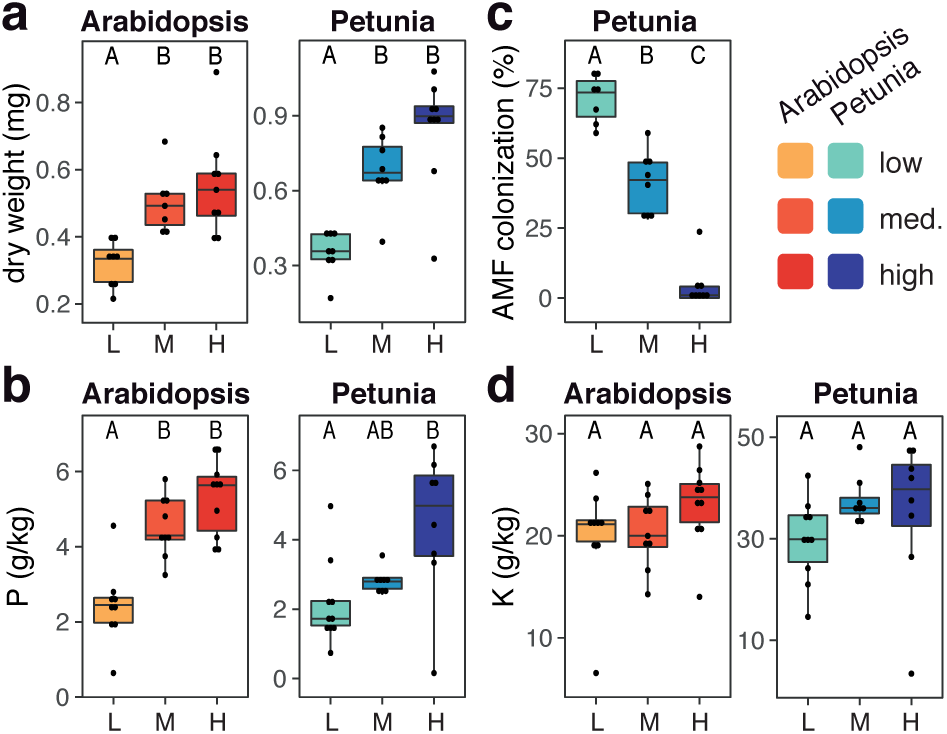
Effectiveness of manipulated phosphate levels on plant growth, leaf nutrient levels and levels of AMF root colonization. Arabidopsis (reddish colors, see legend) and Petunia (blueish colors) were grown at low (L), medium (M) and high (H, increasing hue) levels of P availability and basic plant parameters were recorded to confirm that the experimental setup. Parameters included (**a**) above-ground plant biomass, (**b**) leaf phosphorus levels, (**c**) Petunia root colonization by arbuscular mycorrhizal fungi (AMF) and (**d**) leaf potassium levels. A linear model was used to test for effects of P availability for panels (**a**) (log-transformed data), (**b**) and (**d**) and a quasibinomial generalized linear model for panel (**c**). Different letters indicate signi?cant pairwise differences among sample groups (P < 0.05, Tukey HSD).

### Profiling Soil and Root Microbial Communities

First, we evaluated the following PCR approaches to profile root fungal communities: ITS1F and ITS2 (McGuire *et al.*, 2013a), fITS7 and ITS4 (Ihrmark *et al.*, 2012a) and ITS1F with the reverse complement of fITS7. Community profiles were inspected for the proportions of plant and AMF sequences as well as for fungal diversity. We selected ITS1F and ITS2, because this PCR approach captured low levels of plant sequences at good coverage of AMF and highest levels of taxa richness (**Fig. S2**, See **Notes S1** for a detailed comparison of the PCR approaches).

We then characterized soil and plant root-associated bacterial and fungal communities by sequencing amplicons of the 16S rRNA gene and the internal transcribed spacer (ITS) region 1, respectively. We obtained 2’196’310 high-quality bacteria sequences with a median of 36’718 sequences per sample and 3’809’350 high-quality fungal sequences with a median of 54’337 sequences per sample. Bacterial and fungal sequences clustered into 3’701 bacterial operational taxonomic units (bOTUs) and 1’688 fungal OTUs (fOTUs), respectively.

Soil bacteria comprised abundant Acidobacteria, Actinobacteria, Firmicutes, Deltaproteobacteria and Verrucomicrobia, whereas plant roots were mainly colonized by Betaproteobacteria, Gammaproteobacteria and Bacteroidetes (**Fig. 2**). The bacteria community composition at Phylum rank was not markedly different between Arabidopsis and Petunia.

**Figure 2:**
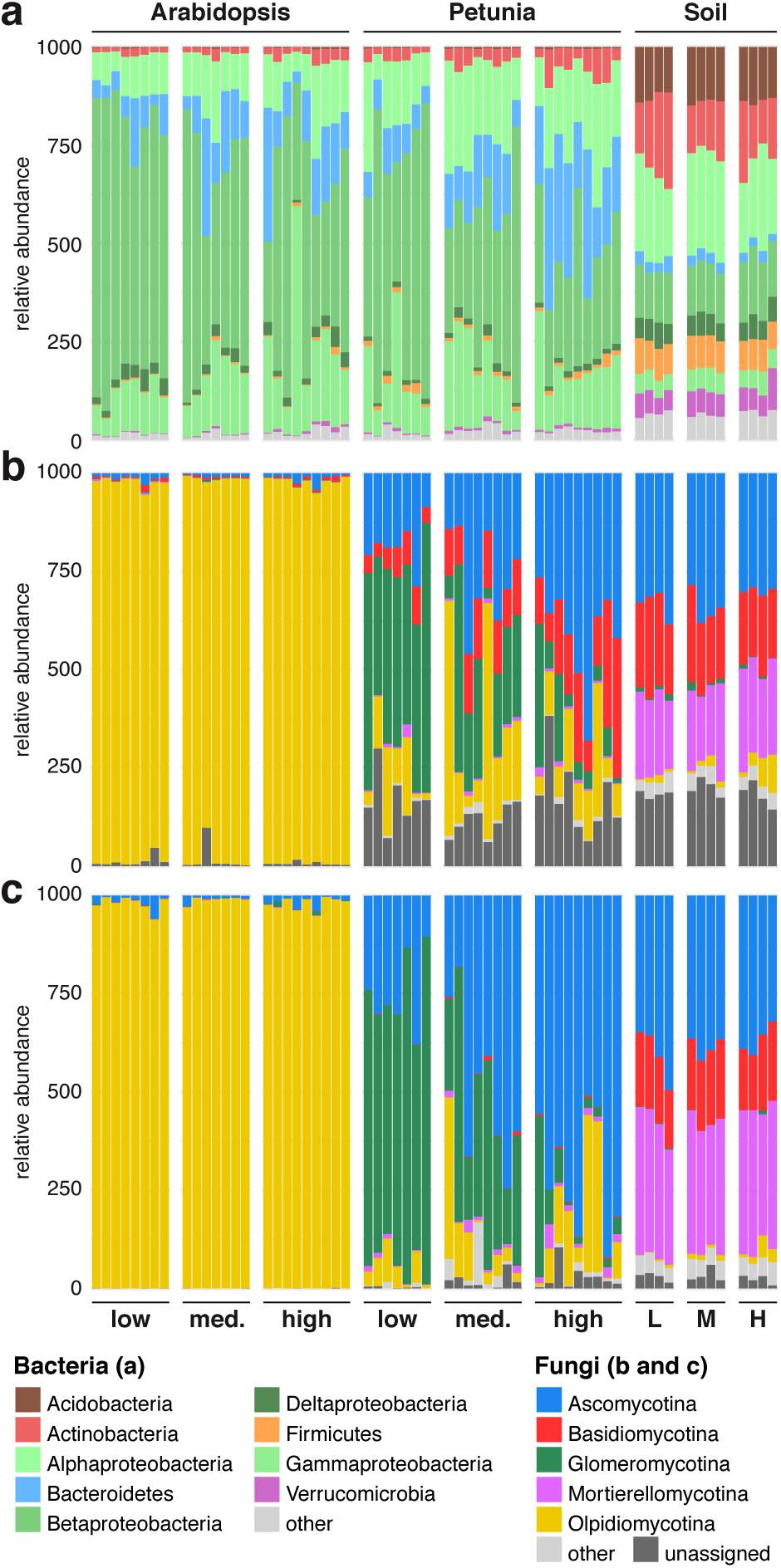
Taxonomic profiles of microbial communities at phylum level. (**a**) Bacteria profiles were obtained using MiSeq sequencing while fungal profiles were determined using (**b**) MiSeq and (**c**) SMRT sequencing. Phyla with relative abundances lower than 1% were summarized with ‘other’. Levels of P-availability are indicated with low (L), medium (med.; M) or high (H).

For fungi, Ascomycota and Basidiomycota were abundant in soil and Petunia root samples (**Fig. 2**). Mortierellomycotina were particularly abundant in soil fungal communities, whereas a high number of Glomeromycotina was found in Petunia roots, which varied as a function of the applied phosphate levels. Consistent with the levels of AMF root colonization measured by microscopy (**Fig. 1c**), Glomeromycotina were most abundant in Petunia roots under low-P conditions and decreased in proportion with increasing P availability (**Fig. 3a**). The cumulative relative abundance of Glomeromycotina sequences was significantly positively correlated (adj. R^2^ = 0.59; *P* < 0.001) with the rate of AMF root colonization as assessed by microscopy (**Fig. 3b**). We noted that the root fungal community of Arabidopsis was dominated by sequences belonging to Olpidiomycotina. Most of these sequences belonged to fOTU1 (assigned to *Olpidium brassicae*), which accounted for 94.50% of the sequences in Arabidopsis but only 0.79% of the sequences in Petunia samples. To exclude that this is a technical peculiarity of MiSeq, we confirmed the dominance of *O. brassicae* by sequencing the entire ITS region (PCR primers ITS1F and ITS4) using SMRT sequencing (**Fig. 2c**). We refer to the **Notes S5** for the detailed comparison of the sequencing approaches. In brief, the SMRT-sequencing based community profiles also avoided amplifying plant sequences while abundantly capturing the AMF. Both methods have their inherent technical advantages with the MiSeq approach offering enhanced throughput and sampling depth, whilst the SMRT-sequencing method provides enhanced taxonomic resolution. Albeit a few quantitative differences, the two approaches reproduce overall similar taxonomic compositions and revealed remarkably similar biological patterns (**Notes S5**).

**Figure 3:**
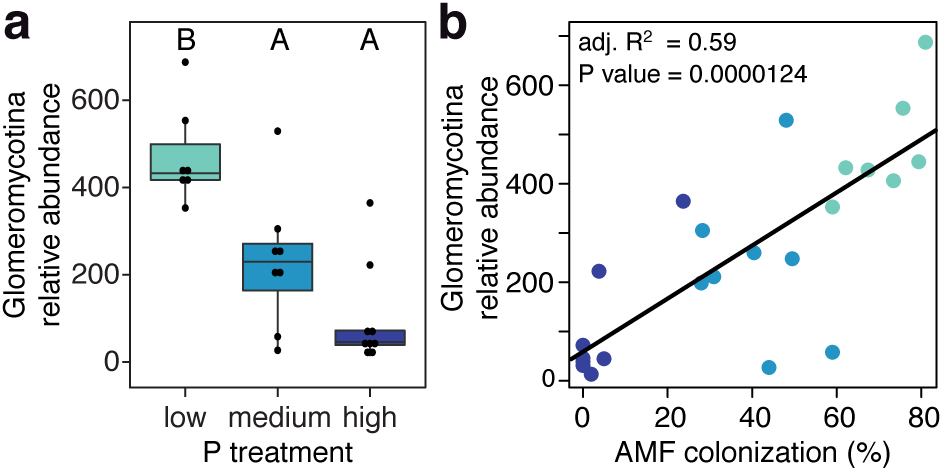
Abundance of AMF in Petunia roots. (**a**) Quantification of AMF based on the relative abundance of Glomeromycotina sequences in the microbiota profiles. Relative abundances were calculated from total sum normalized data. A quasibinomial generalized linear model was used to test for effects due to P availability. Different letters indicate signi?cant pairwise differences between sample groups (P < 0.05, Tukey HSD). This data was correlated (**b**) with the levels of AMF root colonization as measured in the same samples by microscopy (data presented in **Fig. 1c**).

### Phosphate Induced Variation in Microbial Diversity

In the following, we used the MiSeq-based fungi profiles as they were obtained using the same sequencing platform as the bacteria and because of the enhanced sequencing depth. Bacteria and fungi richness was highest in unplanted soil, followed by Petunia and then Arabidopsis roots (**Fig. S3**). ANOVA confirmed the effect of plant species on alpha diversity for both bacteria and fungi and further uncovered an effect by the different P availability on the bacteria community (**Fig. S4**, **Table S1**). Bacterial richness, diversity and evenness were generally higher in Petunia compared to Arabidopsis and generally increased with increasing P concentrations. With the dominance of *O. brassicae*, fungal richness, diversity and evenness were markedly lower in Arabidopsis compared to Petunia.

Utilizing principal coordinate analysis (PCoA) of Bray-Curtis dissimilarities, we found compositional differences in microbial communities due to the tested experimental factors sample type, plant species and P availability (**Fig. 4**). Consistent with previous work (Bulgarelli *et al.*, 2012; Hartman *et al.*, 2018), bacteria and fungi differed markedly between the sample types of unplanted soil and roots (**Fig. S5**). With regard to plant species, fungal communities were more divergent between Arabidopsis and Petunia roots compared to bacteria communities (**Fig. 4**), possibly reflecting their opposite behavior with AMF. Permutational multivariate analysis of variance (PERMANOVA), finding significant plant species effects on both bacterial and fungal communities (**Table S2**), confirmed that plant species explained more variation for fungi (53% of variation) compared to bacteria (14%).

**Figure 4:**
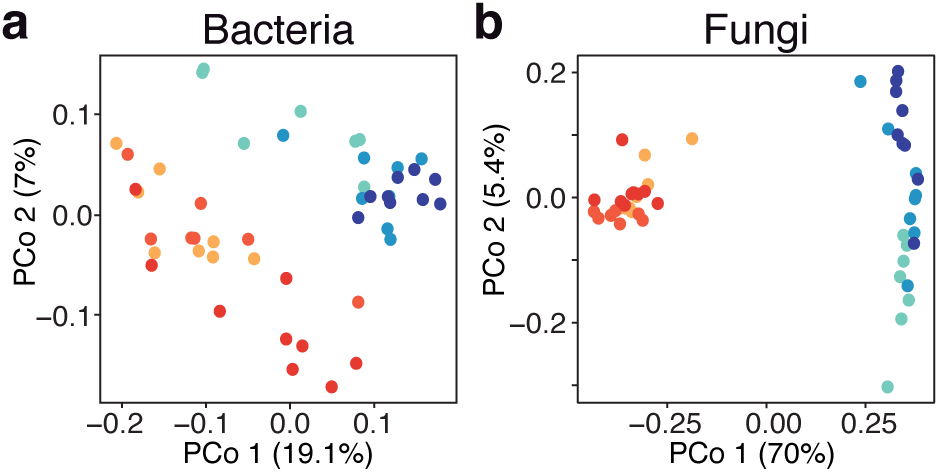
Effects of plant species and P-levels on community composition. Unconstrained ordinations with PCoA using Bray-Curtis dissimilarities were performed on the (**a**) bacterial and (**b**) fungal communities associated with roots. Samples were colored following the color scheme defined in **Fig. 1** (Arabidopsis and Petunia with reddish and blueish colors, respectively, and the increasing P availability (low, medium to high) are marked with increasing hue.

The effects of P availability were apparent in bacterial communities of both Arabidopsis and Petunia by clustering following the gradient in phosphate levels (**Fig. 4A**), whereas for fungi, this was only manifested in Petunia (**Fig. 4B**). PERMANOVA confirmed a significant P availability effect for the bacteria (**Table S2**). To approximate the effect sizes of P availability on the Petunia and Arabidopsis root microbial communities, we inspected the R2 values of PERMANOVA applied to the data of each plant separately. While P availability explained 14.1% and 13.3% of variation in Petunia root bacterial and fungal communities, respectively, it accounted for 15.0% and 21.7% of variation in the Arabidopsis root microbial communities (**Table S3**). This indicates that varying P availability in soil can account for about 15% of variation in plant root microbiota composition.

### Identifying Phosphate Sensitive Microbes

Next, we identified P sensitive OTUs – OTUs being differentially abundant between low and high-P conditions – using edgeR (Robinson *et al.*, 2010). In total we found 2.2% bOTUs and 6.3% fOTUs responsive to the varying P availability in Petunia, while 3.1% bOTUs and 13% fOTUs were sensitive in Arabidopsis (**Fig. 5a**, **Table S4**). With the exception of four bOTUs, different sets of P sensitive bacteria and fungi OTUs were found for Arabidopsis and Petunia, suggesting that the two plant species have differential microbial responses to low-P conditions. Among the four shared bOTUs was a prominent *Dechloromonas* sp. (bOTU2), which is more abundant under low-P conditions in both plant species (**Fig. 5b**). Bacteria from different taxonomic lineages were abundant under low or high-P conditions (**Table S4**). The most abundant Arabidopsis root bacteria included also Burkholderiales, Bdellovibrionales and Rhodocyclales under low-P conditions, whereas taxa from the Chthoniobacterales, Planctomycetales and Verrucomicrobiales were enriched under high-P conditions. Under low-P conditions, the abundant Petunia root bacteria included members of the Burkholderiales and Rhodocyclales, whereas under high-P conditions, a slightly different set of bacteria, including a Flavobacterium sp. (Flavobacteriales), a Tahibacter sp. (Xanthomonadales) and members of the Verrucomicrobiales were abundant. Examples of highly abundant and low-P specific Burkholderiales and Rhodocyclales members include a *Dechloromonas* sp. (bOTU2) and a *Candidatus* Accumulibacter (bOTU13, **Fig. 5b**). Among the Petunia root bacteria, which are enriched under low-P conditions, we noticed an bOTU assigned to *Candidatus* Glomeribacter gigasporarum (bOTU134, **Fig. 5b**), which presents an endobacterium associated with lineages in the AMF family Gigasporaceae (Bianciotto *et al.*, 2003).

**Figure 5:**
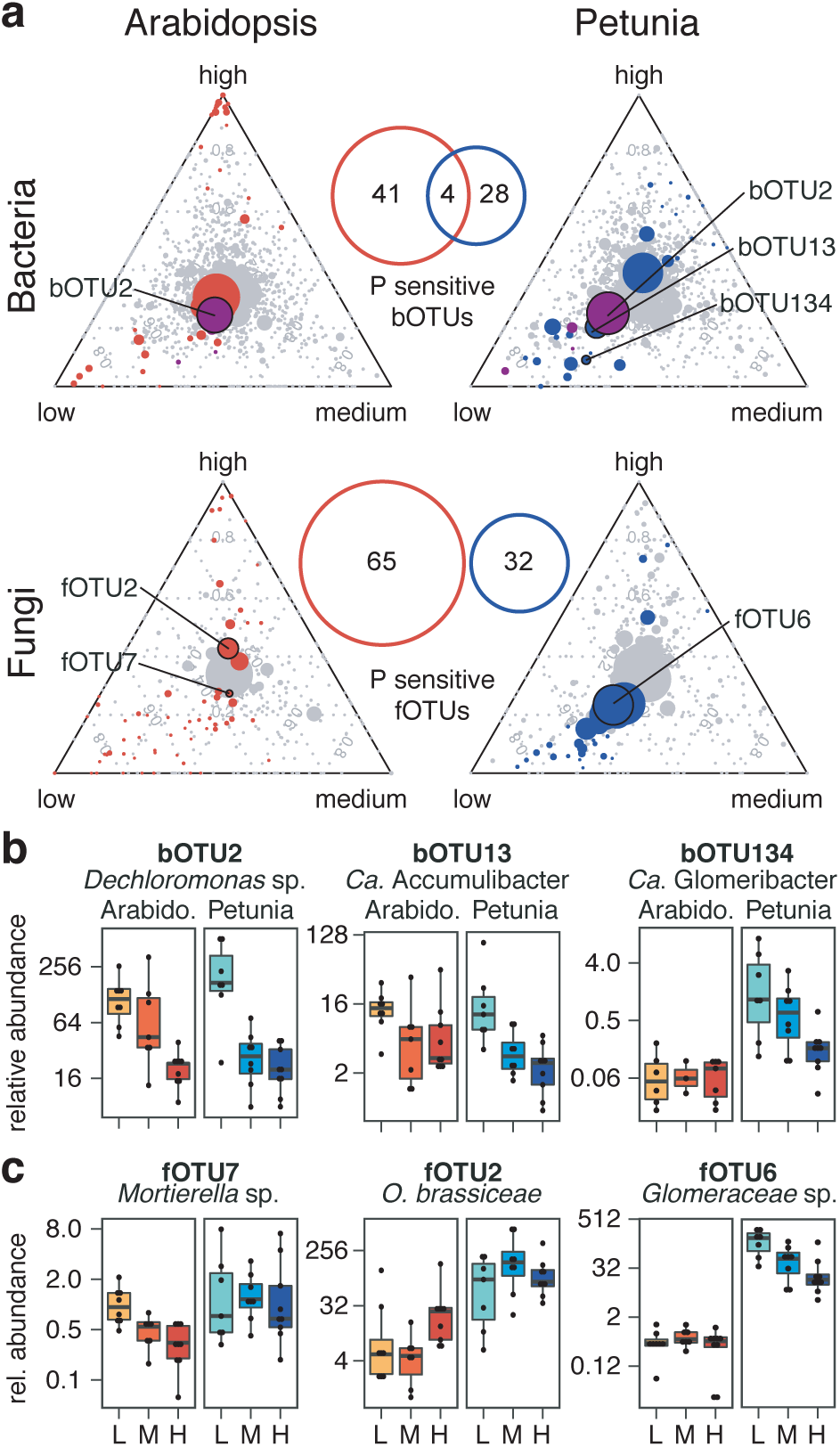
Identifying phosphate sensitive microbes. P sensitive OTUs (differentially abundant between low and high P conditions) were separately identified for the bacteria and fungi, both in Arabidopsis and Petunia (edgeR analysis, high vs low P, FDR < 0.05). (**a**) The ternary plots depict individual OTUs (in circles), sized by their relative abundance, and the position of the OTUs in the triangle reflects their proportional abundance in low, medium and high P samples. P sensitive OTUs are colored in red (Arabidopsis), blue (Petunia) or purple (found in both species) and non-affected OTUs are colored in gray. The number of P sensitive OTUs and their overlap between species is given with the venn diagrams. Panels **b** and **c** illustrate the relative abundances (per milles) of a few representative P sensitive bacterial and fungal OTUs in low (L), medium (M) and high (H) P conditions (They are also indicated in the ternary plots).

Similar to bacteria, different fungal lineages responded to low or high-P conditions in Arabidopsis and Petunia (**Table S4**). In Arabidopsis, besides many low abundant and often taxonomically poorly resolved fungi, the distinct group of Mortierellomycotina (e.g., fOTU7, **Fig. 5c**) was enriched under low-P conditions. Under high-P conditions, the abundant fungi *O. brassiceae* (Olpidiales, fOTU2, **Fig. 5c**), *Hygrophoraceae* sp. (Agaricales, fOTU10) and *Cadophora* sp. (Helotiales, fOTU14) were found besides numerous low abundant fOTUs. While in Petunia only a handful of diverse and low abundant fungi were enriched under high-P conditions, we found a large group of 28 mycorrhizal fOTUs enriched in the low-P treatment (**Table S4**). These mycorrhizal fOTUs belonged mostly to the order Glomerales and included numerous abundant members such as *Funneliformis* and *Glomus* spp. (e.g., fOTU6 in **Fig. 5c**).

### Phosphate-Induced Dynamics in Microbial Abundance

Finally, we utilized co-occurrence network analysis to find pairs or groups of microbes with a similar abundance behavior along the gradient of plant-available P. Co-abundance presents a pre-requisite for cooperation among microbes and we speculated to identify possible candidate cooperation partners that may contribute to support plant growth under low-P conditions. **Figure 6a** visualizes the significant positive pairwise correlations between root microbiota members (bOTU-bOTU, fOTU-fOTU and bOTU-fOTU) of Petunia and Arabidopsis growing in conditions with low, medium or high-P availability. We then partitioned the network into discrete community modules and mapped the P-sensitive bOTUs and fOTUs into the network and modules. While we did not find groups of co-occurring OTUs (=modules) that were responsive to high-P conditions, we found two major modules, ‘M1’ and ‘M26’, that comprised high proportions of P-responsive OTUs (**Fig. 6b**) being specifically abundant under low-P conditions (**Fig. 6c**). The module ‘M1’ comprised only bacteria, mainly belonging to the Betaproteobacteria orders Burkholderiales and Rhodocyclales (**Table S5**). In contrast, the module ‘M26’ grouped a set of five taxonomically diverse bacteria lineages with a large set of fOTUs primarily belonging to the order Glomerales (**Table S5**). These fOTUs represented almost all AMF fOTUs in the dataset (**Table S5**) and interestingly, they co-occurred with the *Candidatus* Glomeribacter bOTU134.

**Figure 6:**
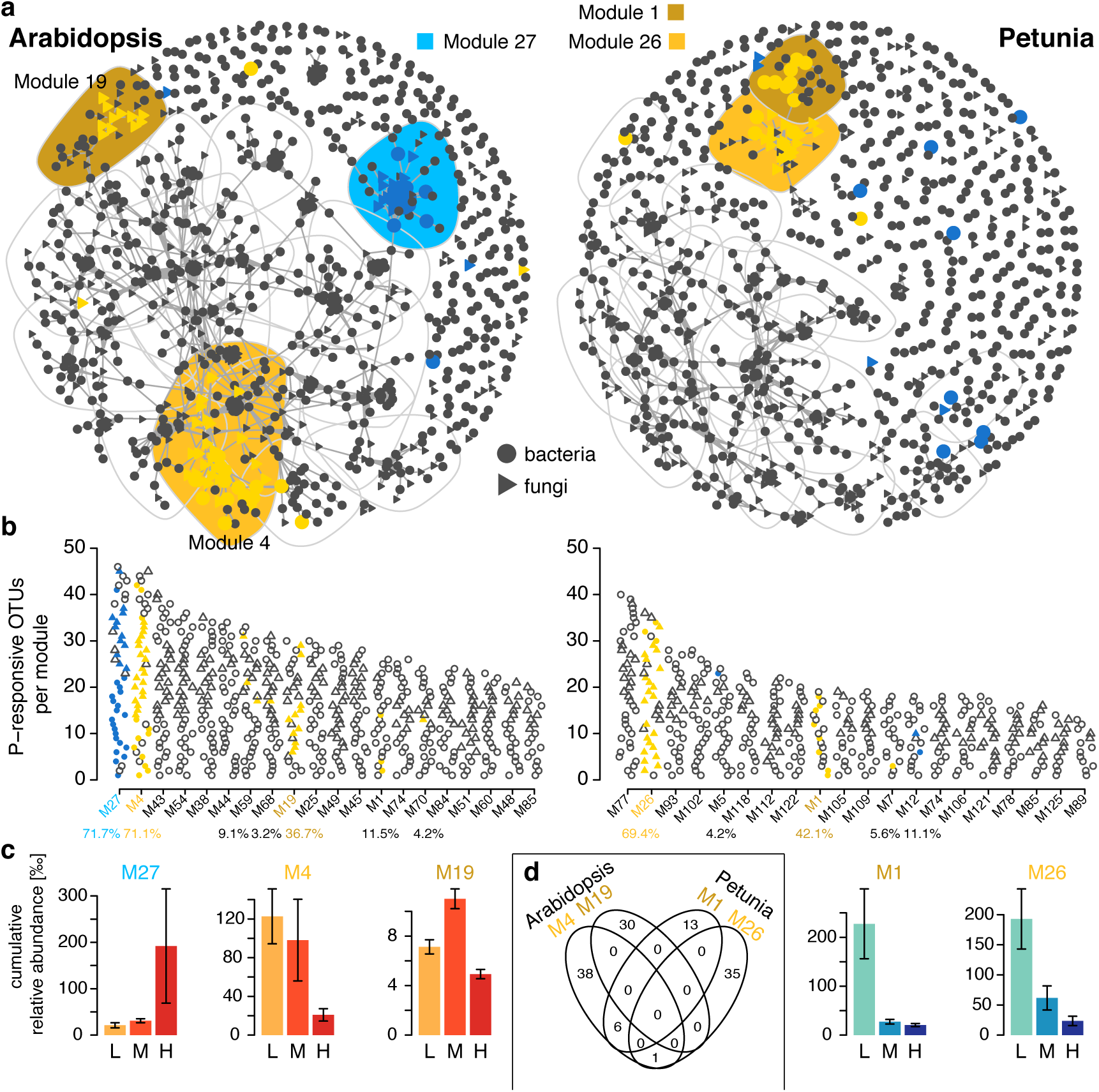
Microbial co-occurrence patterns along the gradient of plant-available phosphate. (**a**) Co-occurrence networks visualize the significant positive pairwise correlations (ρ > 0.7, *P* < 0.001; indicated by links between OTUs) between bacteria (circles) and fungi (triangles) OTUs in Arabidopsis and Petunia root communities. P sensitive OTUs, which are abundant under low and high P conditions, are colored in yellow and blue, respectively. The twenty network modules comprising highest numbers of OTUs are rimmed with grey lines with the modules containing high proportions of P-responsive OTUs being shaded in yellow and blue. (**b**) Top twenty most populated modules, ranked by decreasing numbers of OTUs (bOTUs in circles; fOTUs as triangles) with low and high P sensitive OTUs being colored in yellow and blue, respectively. Percentages below the x-axis report the proportion of P sensitive OTUs present in each module. (**c**) Cumulative relative abundance (as permilles) of all bacteria and fungi OTUs in the P sensitive modules in low (L), medium (M) and high (H) P conditions. The cumulative relative abundance indicates the overall response of the microbes in the P sensitive modules. (**d**) Number and overlap of OTUs in the low P sensitive modules of Arabidopsis and Petunia are shown with the venn diagram.

The same analysis was conducted for Arabidopsis and revealed a module ‘M27’ with co-occurring bacteria and fungi OTUs that were specifically abundant under high-P conditions (**Fig. 6**, **Table S5**). This module grouped diverse bacteria members including Planctomycetes and Verrucomicrobia and a diverse set of fungi. The module ‘M19’ held low abundant and taxonomically diverse bacteria and fungi that favored intermediate P-levels. The module ‘M4’ comprised abundantly co-occurring bacteria and fungi under low-P conditions, belonging mainly to diverse Proteobacteria and Ascomycota or unknown fungi, respectively. With the exception of a few bacteria, the low-P responsive modules of Arabidopsis and Petunia had specific compositions (**Fig. 6d**), which is consistent with the species-specific root microbiota dynamics to low-P condition.

Microbes that simultaneously co-occur with many others are often referred to keystone taxa as they may play an important ecological role by determining community dynamics and microbiome functioning (Banerjee *et al.*, 2018). We identified keystone OTUs, defined based on their high degree of co-occurrence, for the Arabidopsis and Petunia root microbiota networks (**Table S5**). While all Arabidopsis keystone OTUs belonged to the high-P module ‘M27’, we found keystone OTUs in the low-P responsive module ‘M26’ of Petunia. These low-P responsive keystone OTUs were the Glomeromycotina fOTU6 and fOTU111 as well as fOTU109 of unknown taxonomy.

### Endobacteria

To understand whether bacterial communities associated with the Arabidopsis or Petunia root microbiota could include fungal endobacteria, we aligned candidate bOTU sequences to a database of curated endobacteria sequences (see methods). Phylogenetic placement confirmed bOTU134 as *Candidatus* Glomeribacter closely related to one hosted in *Scutellospora pellucida* (**Figs. 7a, S6, S7**). In addition, we identified two bOTUs (330 and 778) mapping to *Mycoplasma*-related endobacteria identified in the AMF species *Claroideoglomus claroideum* and *C. etunicatum*, respectively. Although these two bOTUs do not belong to the ‘AMF module M26’ of Petunia, they were, similar to bOTU134, significantly higher in abundance under low-P conditions (**Table S3**). Phylogenetic placement analysis revealed six additional *Burkholderia*- and two *Mycoplasma*-related endobacteria OTUs (**Figs. 7, S6**), however as they were detected with only a handful of reads in a few samples, we did not include them for network analysis (see **Notes S4** for details). The use of microbiota network characteristics was generally not indicative for identifying endobacteria OTUs, by contrast the clustering-based approach proved to function well (**Figs. S6, S7**, **Notes S4**). In summary, the combined sequencing of bacteria and fungi permitted to identify three endobacteria OTUs that had a consistent abundance behavior with their mycorrhizal hosts along the P-gradient.

**Figure 7:**
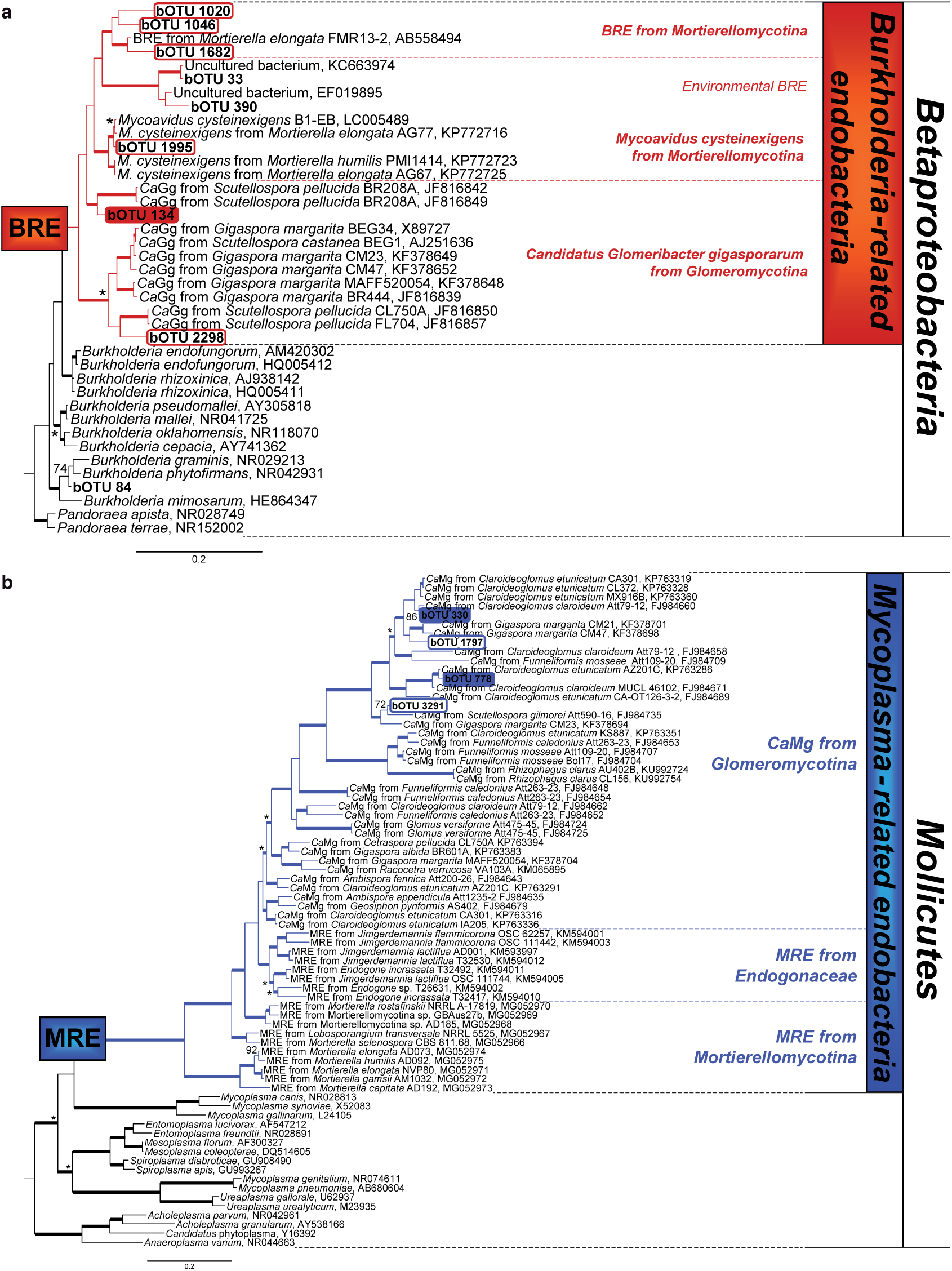
Phylogenetic placement of endobacteria OTUs from candidates identified by the clustering approach. Simplified trees summarizing the confirmed Burkholderia-related endobacteria (BRE, **a**) and Mollicutes-related endobacteria (MRE, **b**); the detailed tree is available as **Fig. S6**. **(a)** Four candidate BRE OTUs cluster within two clades encompassing BRE sequences from Mortierellomycotina fungi. In detail, bOTU 1995 is sister to the type strain of *Mycoavidus cysteinexigens*, whereas bOTUs 1020, 1046 and 1682 cluster with an undescribed BRE hosted in *Mortierella elongata*. Two bOTUs (134 and 2298) cluster within two clades encompassing *Candidatus* Glomeribacter gigasporarum (*Ca*Gg) sequences retrieved from *Scutellospora pellucida* (Glomeromycotina). Two bOTUs (33 and 390) cluster within a new BRE clade, together with putative environmental BRE sequences. **(b)** Four candidate MRE OTUs cluster within different clades encompassing *Candidatus* Moeniiplasma glomeromycotorum (*Ca*Mg) sequences from Glomeromycotina fungi. In detail, bOTUs 330 and 778 cluster with *Ca*Mg hosted in several strains of Claroideoglomus spp., whereas bOTUs 1797 and 3291 cluster with *Ca*Mg hosted in several strains of *Gigaspora margarita* and *Scutellospora pellucida*. The trees show the topology obtained with the Bayesian method. Branches with Bayesian posterior probabilities (BPP) ≥0.95 and ML bootstrap support values ≥70 are thickened; asterisks (*) indicate branches with BPP ≥0.95 but ML bootstrap support values <70; ML bootstrap support values ≥70 are shown for branches having BPP <0.95. Sequences generated in this study are in bold.

## Discussion

In this study, we revealed the dynamics of the root microbiota of non-mycorrhizal Arabidopsis and mycorrhizal Petunia plants along a gradient of plant-available P in soil. We demonstrate that the composition of the root microbiota alters depending on P availability and we revealed species-specific microbial patterns in response to low-P conditions. Under low-P conditions, we confirmed a substantial colonization by AMF in Petunia together with numerous bacteria of the Burkholderiales and Rhodocyclales, whereas Arabidopsis roots hosted mainly Mortierellomycotina fungi and abundant bacteria from the Burkholderiales, Bdellovibrionales and Rhodocyclales. However, the root microbiota of the two plants contained different members (bOTUs) of these taxonomic lineages. These groups of P-sensitive microbes responded simultaneously to the different levels of P availability. Among these co-occurring microbes, we found fungal endobacteria and their corresponding hosts, presenting a well-known example of highly specific multitrophic microbial interactions.

### Endobacteria

To our knowledge, this is the first report to identify endobacteria of Mucoromycotina in a plant microbiota study. Above all, the experimental design with low- to high-P conditions was instrumental to become aware of fungal endobacteria in the dataset. A first hint for the presence of endobacteria came from the statistical approach to find differentially abundant bOTUs between low and high-P conditions, revealing the enrichment of a *Candidatus* Glomeribacter OTU under low-P conditions (**Fig. 5b**). Of note, bOTU134 was almost overlooked, as a reliable taxonomy assignment (confidence >0.7) was only available down to family level, while the deeper data exploration indicated a link with endobacteria (genus-level assignment was ‘*Candidatus* Glomeribacter’, confidence 0.34). The second hint that pointed to endobacteria and their hosts were the co-abundance patterns, which grouped bOTUs including *Candidatus* Glomeribacter bOTU134 with numerous Glomeromycotina fOTUs (**Table S5**). Here again, experimental design was instrumental because the co-occurrence analysis groups microbes with similar abundances in samples from low- to high-P conditions. Finally, we relied on phylogenetic placement to confirm that bOTU134 represents a Candidatus *Glomeribacter*, which is phylogenetically related to a previously isolated exemplar from the AMF species *Scutellospora pellucida* (**Fig. 7**).

The identification of bOTU134 as a fungal endobacterium prompted us to search for additional potential endobacteria in the Arabidopsis or Petunia root microbiota. We compared two strategies, co-occurrence characteristics and sequence similarity to known endobacteria, for their usefulness to identify endobacteria from root microbiota data. For the co-occurrence characteristics strategy, we selected all bOTUs that significantly co-occurred with fOTUs from lineages known to host endobacteria, and then placed the resulting 129 candidate bOTUs into the endobacteria reference tree. For the second strategy, we clustered all sequences of the curated endobacteria database with all representative bOTU sequences of the microbiota dataset, and then placed the resulting 22 candidate bOTUs with the highest sequence similarity into the endobacteria reference tree. Overall, in addition to bOTU134, two abundant (bOTUs 330 and 778) and 8 low-abundant endobacteria OTUs were identified (**Notes S4**). Moreover, the relative abundance of the three OTUs along the P-gradient was consistent with the one of their fungal hosts (**Table S4**). While both approaches identified bOTUs 134 and 778, the clustering-based approach functioned more efficiently as also low abundant candidates were identified.

There are probably multiple reasons why endobacteria and their hosts were largely neglected in microbiota studies so far. Reasons include the low taxonomic resolution of short-read community data or the underrepresentation of reference endobacteria sequences in commonly used taxonomy databases. Here, only the combined sequencing of bacteria and fungi together with the dedicated experimental design to manipulate the abundance pattern of fungal hosts and the curated endobacteria database permitted to identify endobacteria OTUs. Our study demonstrates that microbiota and/or metagenomic datasets represent useful tools to investigate endobacterial–fungal interactions in their true ecological context. Possibly, such cultivation-independent methods can point to further endobacterial–fungal partnerships. Since endobacteria are ecologically relevant for the fitness of their (plant-associated) fungal hosts (Salvioli *et al.*, 2016; Uehling *et al.*, 2017; Desirò *et al.*, 2018), they may also relay some benefits or detriments to the host plant of the fungus (Bonfante & Desirò, 2017). Hence, mycorrhizal plants, their colonizing fungi along with their endobacteria form an entity of a multi-kingdom symbiosis. As put forward by the holobiont concept (Vandenkoornhuyse *et al.*, 2015), the importance of multi-kingdom microbe-microbe interactions for plant performance is not only true for endobacteria and their fungal hosts but also in general between root bacteria and root fungi. For instance, root bacteria are essential to protect plants against pathogenic root fungi (Durán *et al.*, 2018).

### Quantifying AMF in Plant Roots

Although specific sequencing methods to quantify AMF in plant roots (Öpik *et al.*, 2009; Schlaeppi *et al.*, 2016) are available, we sought to establish a sequencing-based approach to assess AMF in plant roots in the context of the whole fungal diversity. In the search for a PCR approach that avoided amplification of plant ITS sequences, we also wanted that AMF would be well captured in plant roots unlike other plant root-fungi profiling methods (Ihrmark *et al.*, 2012b; Hartman *et al.*, 2018). The Illumina approach by McGuire *et al*. (2013b), although reporting soil fungal profiles, indicated that PCR primers ITS1F and ITS2 would permit to abundantly capture AMF and this approach turned out to be successful on plant roots, too (**Fig. S2**).

Prior to community sequencing, we had evaluated the effectiveness of our experimental P availability gradient by confirming that AMF abundantly colonize Petunia roots under low-P but not under high-P conditions (**Fig. 1c**). We performed this quality control using the traditional ‘magnified intersection’ (McGonigle *et al.*, 1990) microscopy method on equivalent subsamples as the ones that were used for the sequencing. The microscopy method relies on staining cleared roots, interpreting and enumerating the different fungal structures following a defined counting scheme under the microscope. Compared to the sequencing-based quantification of AMF in plant roots, the microscopy method appears disadvantageous as it is prone to operator-to-operator variation, is time consuming and lacks throughput, discrimination between AMF species as well as the context of the whole fungal diversity. Nevertheless, we found a reliable agreement between the two methods with a significant positive correlation (**Fig. 3b**). In summary, the MiSeq-based community profiling approach with the PCR primers ITS1F and ITS2 avoided amplification of plant ITS sequences, abundantly displayed the AMF and independently reproduces AMF colonization patterns of plant roots.

### AMF as Keystone Species

Keystone taxa are thought to own central positions in microbial networks, with many links to other species; therefore, they may play important ecological roles by determining community dynamics and microbiome functioning (Banerjee *et al.*, 2018). AMF were postulated to be keystone species (van der Heijden & Hartmann, 2016) and indeed, we find two Glomeromycotina OTUs (fOTU6 and fOTU111) as being keystones in the low-P responsive Petunia module ‘M26’ (**Fig. 6**). Technically, the two Glomeromycotina OTUs fulfilled the network-based criteria defining keystone OTUs – keystone OTUs have high degree of co-occurrence, high closeness centrality and low betweenness centrality (Banerjee *et al.*, 2018). The two identified keystone OTUs have high degrees of co-occurrence implying they are co-abundant with many other (mostly Glomeromycotina) OTUs in the network. However, with regard to these network-topology based criteria, there is a major caveat linked to Glomeromycotina fungi: several Glomeromycotina species have particularly high intraspecies genetic diversity at the rRNA operon (Stockinger *et al.*, 2010; Lekberg *et al.*, 2014) and, as a consequence, these fungi require multiple OTUs (at >97% sequence identity) to represent a single species. For example, three OTUs described the AMF *Rhizoglomus irregulare* (Schlaeppi *et al.*, 2016). Therefore, multiple OTUs per fungal species artificially inflate the number of co-occurring OTUs for an AMF species (the 3 OTUs of *R. irregulare* will consistently co-occur with each other). While we agree that AMF fulfil a key role in root microbiome functioning and this is especially true under low-P conditions, we anticipate that the mathematical criteria for identifying AMF keystones from network data may need to be improved and validated using functional analyses. Instead of assigning keystone status to some pseudo-replicated sequence groups (e.g., the 3 OTUs for *R. irregulare*) based on network topology, we rather favor the idea of empirically nominate keystone microbes because of their known key function(s) in a given condition (e.g., low-P availability).

### Olpidium Brassicae

The root microbiota of Arabidopsis but not Petunia, whether profiled with MiSeq or SMRT-sequencing, comprised abundant sequences assigned to *O. brassicae*, which is a common root-infecting fungal pathogen of *Brassicaceae* plants (Lay *et al.*, 2018). As biotrophic fungus, *O. brassicae* does not cause tissue maceration. Although the Arabidopsis roots were without signs of disease when harvested, we learned after the community sequencing that the fungus had spread in the root tissue. Nevertheless, such dominance of *O. brassicae* OTUs was observed in previous studies (Tkacz *et al.*, 2015; Durán *et al.*, 2018; Lay *et al.*, 2018). Similar to our study, Lay *et al*. (2018) examining canola, wheat and pea roots, also found a Brassicaceae-specific enrichment of an *O. brassicae* OTU. Because these findings originate from different Brassicaceae species and different soil types, we consider the dominance of *O. brassicae* in Arabidopsis roots in our study rather a true biological observation than a technical artifact. Although, we confirmed this observation with SMRT sequencing, we cannot fully exclude that the frequent amplification of *O. brassicae* OTUs may be linked to the PCR primer ITS1f, as all these studies target the ITS1 region and have the use of ITS1 or ITS1f PCR primers in common.

### P-Sensitive Microbiota

We found species-specific microbial patterns in response to low-P conditions consisting of abundant Mortierellomycotina and Glomeromycotina fungi in Arabidopsis and Petunia, respectively (**Fig. 5, Table S4**). Similar to the functioning of AMF in plant P provision, there are some reports that *Mortierella* spp. support P nutrition of plants (Alori *et al.*, 2017). Although represented by different sequence groups, bacteria of the Burkholderiales and Rhodocyclales were abundant in both species. Bacteria of both orders, including members that attach to AMF hyphae, are well known for their ability to solubilize and mobilize P (Sharma *et al.*, 2013; Taktek *et al.*, 2015; Alori *et al.*, 2017). *Candidatus* Accumulibacter being abundant in Petunia under low-P conditions (bOTU13, **Fig. 5b**) as well as the *Dechloromonas* sp. (bOTU2, enriched in low-P, both plant species) are both intriguing root bacteria, as they are capable of polyphosphate metabolism, which has been implicated in stress response to low nutrients in the environment (Rao & Kornberg, 1996; Flowers *et al.*, 2013). In summary, it appears that non-mycorrhizal Arabidopsis and mycorrhizal Petunia rely on different microbial associations to cope with low-P conditions.

We revealed species-specific and in contrast to Robbins *et al*. (2018), we found marked root microbiota dynamics of the two plants along a gradient of plant-available. A plausible reason for stronger P effects on the root microbiota is that our approach created stronger P limiting conditions (1.05 mg P kg^-1^ soil) compared to the low-P control soil (P1, did not receive phosphate amendments over the last 65 years) of the long-term P fertilization experiment studied by Robbins et al. (2.3 mg P kg^-1^ soil, values are directly comparable as they were analyzed in the same professional soil laboratory with the same method). Moreover, we established a steeper gradient between low and high-P conditions with a P availability of ∼113 mg P kg^-1^ soil as high-P condition compared to ∼12 mg P kg^-1^ by Robbins *et al*. (2018). The stronger P limiting conditions in our study were also reflected in the different rosette biomass data in both studies as we find a twofold reduction in median rosette biomass whereas they measured at maximum a 1.5x effect in low-P soils.

### Concluding Remarks

The analysis of root microbiota dynamics of Arabidopsis and Petunia to low-P conditions revealed a number of plant-species specific root microbes that are preferentially selected at low soil P availability. With regard to agricultural applications, this works suggests that for supporting different plant species in P nutrition, different P-solubilizing and/or mineralizing bacteria strains are needed. Possibly this explains the high context dependency of successful field applications with P-solubilizing and/or mineralizing bacterial products.

## Acknowledgements

We acknowledge A. Held (Agroscope) for support in amplicon library preparation. We thank Prof. D. Reinhardt from the University of Fribourg for Petunia seeds. We thank Drs. M. Zuber and D. Bürge from Agroscope for the analysis of the leaf nutrient content and soil chemical properties. We thank Drs. L. Poveda and A. Patrignani from the Functional Genomics Center in Zürich for technical support in MiSeq- and SMRT-sequencing, respectively. We acknowledge input for edgeR analysis from Dr. C. Soneson and Prof. M. Robinson from the University of Zürich. This study was supported by the Swiss State Secretariat for Education, Research and Innovation with a project C14.0132 granted to KS and MvdH.

## Author Contributions

NB, MvdH and KS designed the experiments. NB performed the experiment, did the molecular work and analyzed the data. JCW, VS, AD and KS contributed to data analysis. NB, LB, MvdH and KS wrote the manuscript. All authors approved the final manuscript.

This article comprises the following Supporting information:

- Figure S1: Analysis steps

- Figure S2: Comparison of ITS PCR approaches for plant root samples

- Figure S3: Rarefaction curves for bacterial and fungal OTU richness

- Figure S4: Effects of plant species and P-levels on microbial richness, diversity and evenness

- Figure S5: Beta-diversity analysis including the soil samples

- Figure S6: Identification of endobacteria by phylogenetic placement

- Table S1: Effects of plant species and P treatment on alpha diversity (ANOVA)

- Table S2: Effects of plant species and P treatment on community composition (PERMANOVA)

- Table S3: Effects P treatment on species-specific community compositions (PERMANOVA)

- Table S4: Statistics from identifying phosphate sensitive microbes

- Table S5: Network characteristics

- Methods S1: Microbiota profiling and analysis

- Notes S1: Comparison of PCR approaches

- Notes S2: Bioinformatic scripts

- Notes S3: Data analysis in R

- Notes S4: Mapping endobacteria

- Notes S5: Comparison of ITS profiling approaches

